# Acute PFOS exposure consistently dysregulates glucose-stimulated insulin secretion across model systems

**DOI:** 10.1101/2025.10.23.684149

**Authors:** Jana Palaniyandi, Myriam P. Hoyeck, Erin van Zyl, Lahari Basu, Ma. Enrica Angela Ching, Amanda Ameyaa-Sakyi, Amélie Blais, Azam Tayabali, Amy A. Rand, Jennifer E. Bruin

## Abstract

**Aims/hypothesis:** Poly- and perfluoroalkyl substances (PFAS) are fluorinated chemicals widely used in consumer and industrial products. Serum PFAS concentrations have been linked to increased type 2 diabetes risk, but whether this is caused by direct effects of PFAS on the endocrine pancreas remains unclear. This study expands on previous biodistribution data to better characterize PFAS accumulation in human pancreas. We also assessed the effects of perfluorooctane sulfonic acid (PFOS) on pancreatic beta cell function using various model systems.

**Methods:** We measured concentrations of three legacy PFAS (PFOS, PFOA, PFHxS) in plasma and pancreas from 88 human donors. We also modeled PFOS and PFOA exposure in mice to confirm our human biodistribution data. We next exposed immortalized INS-1 cells, primary human donor islets, and primary mouse islets to DMSO (vehicle) or PFOS (1, 10 or 100 μM) either acutely during a glucose-stimulated insulin secretion (GSIS) assay or for 48h (prolonged) and GSIS was subsequently measured. We also assessed whether the acute effects of PFOS on GSIS in mouse islets were mediated by G-protein coupled receptor-40 (GPR40) and/or glucagon-like peptide-1 receptor (GLP-1R). Lastly, we exposed female mice to DMSO or 10 mg/kg PFOS with or without a 2 g/kg glucose bolus to assess the acute effects of PFOS on glucose homeostasis *in vivo*.

**Results:** PFOS was detected at higher concentrations in both plasma and pancreas compared to PFOA and PFHxS in our human donor population. Similarly, PFOS accumulated in pancreas at concentrations 2x higher than PFOA in our mouse model. We provide compelling evidence that PFOS exposure dysregulates insulin secretion in various model systems. Prolonged PFOS exposure suppressed GSIS in INS-1 832/3 cells and mouse islets, whereas acute PFOS exposure stimulated GSIS in mouse islets and in a subset of our human donor islets. We also show that the acute effects of PFOS on GSIS in mouse islets were partly mediated by GLP-1R. Importantly, acute PFOS exposure increased plasma insulin levels in female mice in *vivo*, but only when PFOS exposure was concurrent with a glucose bolus.

**Conclusions:** Legacy PFAS chemicals are consistently detected in human pancreas tissue. We clearly show that PFOS exposure dysregulates insulin secretion in various model systems, although effects varied based on the model and duration of exposure. Collectively, our findings support emerging evidence that PFOS contribute to beta cell dysfunction and points to the importance of model selection when assessing toxicological effects of PFAS on islet function.

## 1. INTRODUCTION

Type 2 diabetes incidence has increased dramatically over the last few decades and is projected to affect over 850 million adults worldwide by 2050 [1]. Dysregulated insulin secretion from pancreatic beta cells— located in the islets of Langerhans—is an early marker of type 2 diabetes progression [2]. Specifically, hyposecretion of insulin will result in hyperglycemia, which can develop into overt type 2 diabetes if left untreated, whereas hypersecretion can lead to insulin resistance and ultimately, beta cell exhaustion [3]. While genetics and lifestyle are known to influence type 2 diabetes risk, these factors alone cannot explain the rapid rise in type 2 diabetes incidence. In fact, there is mounting evidence showing that exposure to persistent organic pollutants (POPs) increases type 2 diabetes risk (reviewed in [4,5]), but their role as beta cell toxicants warrants further investigation.

Poly- and perfluoroalkyl substances (PFAS, aka “forever chemicals”) are characterized by a non-polar fluorinated tail and polar head, making them amphipathic and extremely stable. Given their unique physical-chemical properties, PFAS are used in a wide variety of consumer and industrial applications, including personal care products, fire-fighting foams, and food packaging [6]. The most well-known legacy PFAS analytes include perfluorooctanoic acid (PFOA), perfluorooctane sulfonic acid (PFOS), and perfluorohexanesulfonic acid (PFHxS). Due to their environmental persistence and adverse health effects, there have been extensive regulatory efforts to reduce the global production and use of these long-chain legacy PFAS chemicals [7,8]. This has led to the recent production of alternative “emerging” short-chain analytes [9]. However, PFOS and PFOA continue to be widely detected globally [10,11], with detection rates of >99% in the general Canadian population [12].

Epidemiological studies report a correlation between serum legacy PFAS concentrations and increased risk of pre-type 2 diabetes [13], type 2 diabetes [14,15], and associated metabolic dysfunction, including insulin resistance [16,17] and glucose intolerance [18]. Serum PFAS have also been associated with markers of beta cell dysfunction in humans, including increased fasting insulin [17] and homeostatic model assessment of beta cell function (HOMA-β) [16,17]. It remains unclear whether these adverse metabolic effects are caused by direct action of PFAS on the endocrine pancreas.

To our knowledge, only one study has measured PFOA and PFOS in human pancreas; both compounds were detected in pancreas tissue pooled from 7 human donors, but PFOS was measured at 2-fold higher concentration than PFOA [19]. These data suggest that PFAS can accumulate within the pancreas and may directly impact beta cell health and function. However, given the limited sample size in this study, there is a need for measuring PFAS accumulation in pancreas across a larger cohort of human donors to capture donor-to-donor heterogeneity.

Limited studies have assessed the effect of PFOS on beta cell function, but collectively these studies suggest that PFOS dysregulates insulin secretion [20–23]. Specifically, male, but not female, mice that received an oral gavage of 10 mg/kg PFOS had increased plasma insulin levels 1-hr post-exposure compared to the vehicle control [21]. In line with these data, a 1-hr exposure of an immortalized beta cell line (beta-TC-6 cells) to 50–200 µM PFOS concurrently with 1.4 mM low glucose (equivalent to fasting glucose levels) [21] or 16.7 mM high glucose [20] increased insulin secretion compared to control. However, it is important to note that Zhang et al. [20] did not assess insulin secretion under low glucose conditions prior to stimulating with high glucose, but rather assessed low and high glucose in separate studies; this makes it challenging to assess the effects of PFOS on glucose-stimulated insulin secretion (GSIS) and to calculate stimulation index. Interestingly, both studies in beta-TC-6 cells suggest that G-protein coupled receptor-40 (GPR40) may be partially mediating the stimulatory effects of PFOS on insulin secretion [20,21]; GPR40 is a cell surface receptor on the beta cells that typically binds long-chain fatty acids to stimulate GSIS. Specifically, the effect of PFOS on insulin secretion under low glucose conditions was reduced by ∼50% when beta-TC-6 cells were co-exposed to PFOS and a GPR40 antagonist (GW1100) [21]. These data suggest that PFOS exposure under low or high glucose conditions can acutely stimulate insulin secretion, possibly through GPR40. More detailed studies assessing the acute effects of PFOS on beta cell function are warranted.

Prolonged 48-hr exposure of beta-TC-6 cells to 10–400 µM PFOS significantly suppressed insulin secretion when cells were subsequently incubated with 1.4 mM glucose [22] or 16.7 mM glucose [24]. Similarly, MIN6 cells exposed to 100 pM, 100 nM or 10 µM PFOS for 24-hrs had suppressed insulin secretion when subsequently stimulated with 16.7 mM glucose [25]. Interestingly, the effect of prolonged PFOS exposure on GSIS differed in EndoC-βH1 cells, an immortalized human beta cell line [25]. EndoC-βH1 cells exposed to 10 nM and 1 µM PFOS for 72-hrs had increased basal insulin secretion under 2.8 mM glucose, whereas cells exposed to 10 nM and 100 nM PFOS had increased insulin secretion under 20 mM high glucose conditions [25]. These data suggest that prolonged PFOS exposure suppressed insulin secretion in immortalized rodent cell lines but increases insulin secretion in human cell lines. More work is needed to help understand the discrepancies between cell models.

While immortalized beta cells are a robust and typically glucose-responsive model [26], they lack key features of pancreatic islet morphology and physiology [27,28]. Immortalized cells are highly proliferative and cultured in a 2D monolayer, whereas islets have limited proliferative capacity and are cultured in 3D, allowing for preservation of islet architecture. Another limitation for immortalized beta cells is the absence of other endocrine cells (i.e. alpha, delta, and PP cells) and consequently intra-islet paracrine signaling, which is crucial for the modulation and amplification of GSIS [29]. For example, the alpha cell secretes glucagon and glucagon-like peptide-1 (GLP-1), which binds locally to GLP-1 receptors (GLP-1R) on beta cells and amplifies second-phase GSIS. Delta cells are responsible for secreting somatostatin, which inhibits the release of other endocrine hormones, including insulin and glucagon [30]. Together, these hormones work to control glucose homeostasis. As such, it is important to consider the impact of PFAS on GSIS in primary whole islets.

Our study first expands on previous biodistribution data by measuring concentrations of three legacy PFAS (PFOS, PFOA, PFHxS) in pancreas and plasma of 88 human organ donors to determine the extent and range of PFAS accumulation in human pancreas. Next, we modeled human PFOS and PFOA exposure in mice to confirm our findings from the human biodistribution study. We show convincing evidence that PFOS preferentially accumulates in pancreas of both humans and mice compared to PFOA and PFHxS. As such, we next assessed the acute and prolonged effects of PFOS exposure on beta cell function in immortalized INS-1 832/3 cells, primary human donor islets, and primary mouse islets. Lastly, we assessed whether the acute effects of PFOS on insulin secretion *in vitro* could be replicated in an *in vivo* mouse model.

## 2. MATERIALS AND METHODS

### 2.1. PFAS biodistribution in human donor pancreas and plasma

Human donor pancreas and plasma were processed by the Alberta Diabetes Institute IsletCore at the University of Alberta in Edmonton (http://www.isletcore.ca/) with the assistance of the Human Organ Procurement and Exchange (HOPE) program, Trillium Gift of Life Network (TGLN), and other Canadian organ procurement organizations. All donors’ families provided informed consent for the use of pancreatic tissue and plasma in research. Research with human donor tissue was approved by the Human Research Ethics Board at the University of Alberta (Pro00013094) and at Carleton University (#106701).

#### 2.1.1. Plasma and pancreas collected for mass spectrometry

Human donor plasma (n = 79) and pancreas (n = 88) were collected between January 2022 and December 2023 (donor information in **Supplemental Table 1**). For each donor, plasma (∼1 mL) was obtained and stored at -30°C prior to liquid chromatography tandem mass spectrometry (LC-MS/MS) analysis. A pancreas biopsy (0.363–1.240 g) was collected from the same group of donors, flash frozen and subsequently stored at -30°C prior to LC-MS/MS analysis. All donor characteristics are publicly available on the www.humanislets.com website [31].

#### 2.1.2. LC-MS/MS for human donor samples

Measurement of legacy PFAS concentrations in human plasma and pancreas was completed by SGS AXYS (Sidney, BC, Canada) according to AXYS EPA method 1633. SGS AXYS is ISO 17025 accredited by the Canadian Association for Laboratory Accreditation. Specifically, PFOA, PFOS, and PFHxS concentrations were measured using LC-MS/MS in negative electrospray ionization mode. Samples were spiked with isotopically labelled recovery standards (^13^C_4_-PFOA, ^13^C4-PFOS, ^18^O_2_-PFHxS), extracted by solid-phase extraction using a wax cartridge, and eluted using 1% methanolic ammonium hydroxide. Sample concentrations were quantified using isotope dilution quantification and adjusted for percent recovery (**Supplemental Table 2**). An aqueous blank sample and spiked matrix samples were included in the analysis. Sample limits of detection (LOD) were established based on a 3:1 signal to background noise ratio and were determined for each analyte and sample (**Supplemental table 3**). Non-detectable analytes were assigned a value equivalent to ½ LOD using batch-specific LOD values.

### 2.2. PFAS biodistribution in mouse pancreas

#### 2.2.1. Cohort 1 – Measuring PFOS and PFOA in pancreas following exposure by daily oral gavage

This study was conducted by the Mechanistic Studies Division at Health Canada (Ottawa, Canada). Female and male C57BL/6-Elite mice (8–10 weeks old; n = 3/condition) were purchased from Charles River. Mice were exposed to 0.5% Tween-20 (vehicle) diluted in optima water (Fisher Scientific, #W64; Hampton, NH, USA), 4.5 mg/kg/d PFOA (Sigma-Aldrich, #171468; St. Louis, MO, USA), or 4.5 mg/kg/d PFOS (Sigma-Aldrich, #77283) for 28 days via oral gavage (study design schematic in **Figure 2A**). This dose was selected based on existing toxicological data from the Agency for Toxic Substances and Disease Registry (ATSDR); 4.5 mg/kg/d falls within a range expected to produce measurable biological effects while minimizing the risk of adverse outcomes such as weight loss or severe toxicity in repeated exposure studies [32]. Whole pancreas was collected at study endpoint, snap frozen in liquid nitrogen and stored at -80°C for LC-MS/MS (see *Section 2.2.3*). This study was approved by the Health Canada Animal Care Committee (#2022-019).

**Figure 1.**
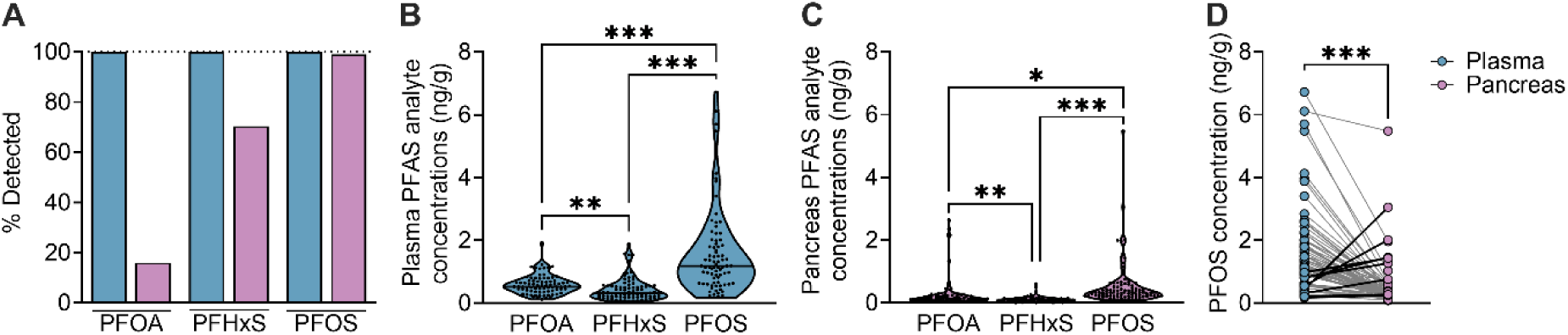
PFOS is consistently detected in human pancreas tissue, and in some donors, at higher concentrations than in plasma. All human organ donor samples were obtained through the ADI IsletCore. Plasma and whole pancreas tissue were collected for mass spectrometry analysis to measure PFOA, PFOS, and PFHxS. **(A)** Percent detection rates for each PFAS analyte measured. **(B,C)** Concentrations of PFOA, PFHxS, and PFOS in human **(B)** plasma (n = 79) and **(C)** pancreas tissue (n = 88). **(D)** A before-after plot comparing PFOS concentrations in human plasma and pancreas for each donor, where data points for a given donor are connected by a line (n = 78). **(B–D)** Individual datapoints represent individual donors. Data is presented as mean ± SEM. *p<0.05, **p<0.01, and ***p<0.001. The following statistical tests were used: **(B,C)** one-way ANOVA with Tukey post-hoc; **(D)** two-tailed paired t-test.

**Figure 2.**
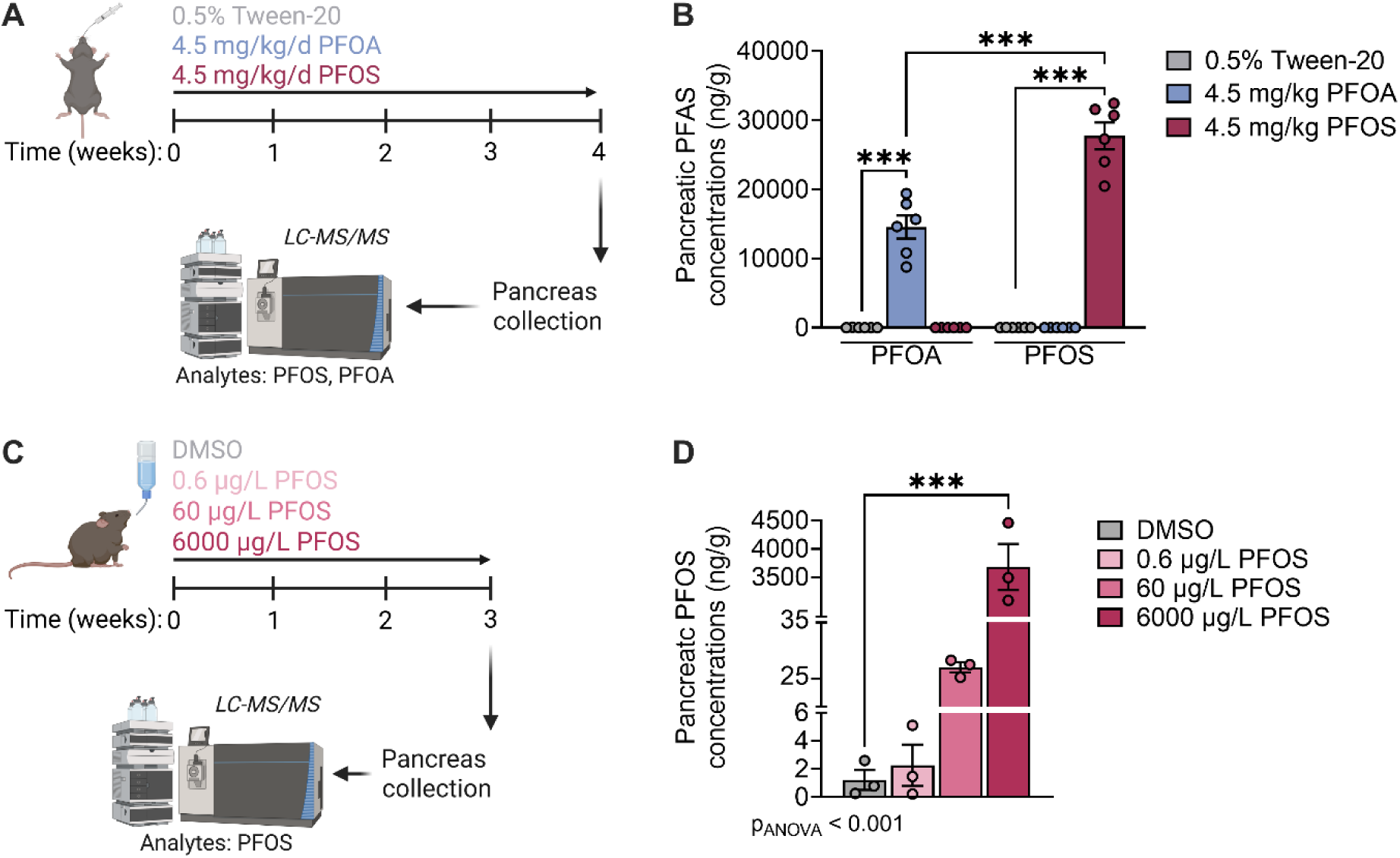
PFOS accumulates in mouse pancreas tissue at higher levels than PFOA. **(A)** Schematic outlining the study timeline for cohort 1. Male and female mice were exposed to 0.5% Tween-20 (vehicle), 4.5 mg/kg/d PFOS, or 4.5 mg/kg/d PFOA via oral gavage for 28 days (n = 6/condition). Pancreas tissue was collected at endpoint for liquid chromatography–tandem mass spectrometry analysis (LC-MS/MS). **(B)** PFOS and PFOA concentrations in mouse pancreas; male and female data was pooled. **(C)** Schematic outlining the study timeline for cohort 2. Female mice were exposed to DMSO (vehicle) or PFOS (0.6, 60 or 6000 µg/L) via drinking water for 21 days. Pancreas tissue was collected at endpoint for LC-MS/MS (n = 3/condition). **(D)** PFOS concentrations in pancreas. Individual datapoints on bar graphs represent biological replicates. ***p<0.001. The following statistical test was used: **(B,D)** one-way ANOVA with Dunnett’s post-hoc.

#### 2.2.2. Cohort 2 – Measuring PFOS in pancreas following daily exposure via drinking water

Female C57Bl/6J mice (8–10 week old; n = 5/condition) were purchased from Jackson Laboratories (Maine, USA) and exposed to 2.9% dimethyl sulfoxide (DMSO; vehicle) or PFOS (0.6, 60 or 6000 µg/L) via drinking water (Milli-Q) for 21 days (study design schematic in **Figure 2C**). Whole pancreas tissue was collected at study endpoint, snap frozen in liquid nitrogen and stored at -80°C for LC-MS/MS (see *Section 2.2.3*). This study was approved by the Carleton University Animal Care Committee and carried out in accordance with the Canadian Council on Animal Care guidelines.

The 0.6 µg/L dose (∼0.00012 mg/kg) corresponds to Health Canada’s maximum acceptable concentration of PFOS in drinking water, a level at which no adverse health effects are expected [33]. We then selected a dose 100x higher than Health Canada’s maximum acceptable concentration (60 µg/L; ∼0.012 mg/kg), which also aligns with PFAS contamination levels in drink water from high-risk populations in the USA (e.g. communities living by fluoropolymer production facilities) [34,35]. The 6000 µg/L dose (∼1.2 mg/kg) is a supraphysiological dose that aligned with the dose used in *2.2.1. Cohort 1*.

#### 2.2.3. LC-MS/MS for mouse pancreas samples

PFOS and PFOA concentrations in mouse pancreas samples (0.073–0.099 g) were measured in Dr. Amy Rand’s lab in the Department of Chemistry at Carleton University (Ottawa, Canada) using a method adapted from Yeung et al [36]. Frozen mouse pancreas samples were homogenized in Milli-Q water, ion paired with basified tetra-*n*-butylammonium hydrogen sulfate and extracted three-times with methyl-tert-butyl ether. Organic layers were separated from aqueous and combined, blown to dryness under a stream of nitrogen gas, and reconstituted in methanol. Prior to LC-MS/MS analysis, particulates were removed from samples with a nylon syringe filter. LC-MS/MS was conducted using a Waters Acquity I-Class UPLC coupled to a Xevo TQS-micro MS/MS. A mixture of ^13^C- and ^18^O-labeled PFAS standards (MPFAC-MXA, Wellington Laboratories) was used for quantification. PFOS and PFOA concentrations were quantified using multiple reaction monitoring (MRM) mode, with relative response ratios calculated against labeled internal standards.

### 2.2. *In vitro* models

#### 2.2.1 PFOS preparation and dosing

PFOS (SynQuest Laboratories, #6164-3-08, Alachua, FL, USA) was solubilized in DMSO (Sigma-Aldrich, #276855) to a final stock concentration of 0.1 M. The solution was kept at -20°C for long-term storage. 0.1% DMSO was used as the vehicle control for all *in vitro* experiments.

Final PFOS exposure concentrations of 1, 10, 100 µM were used. These doses align with previous studies that evaluated the effects of PFOS on insulin secretion *in vitro* [20–22] since we wanted to expand on their work using various model systems. We recognize that the concentrations used exceed plasma concentrations measured in the general population. For example, the 1 µM PFOS dose is ∼170-fold higher than the median PFOS plasma concentration detected in Canadians (3.0 µg/L or 0.0059 µM), and ∼45-fold higher than the 95^th^ percentile of PFOS plasma concentration detected in Canadians (11 µg/L or 0.022 µM) [12]. However, our doses are congruent with high-dose exposure scenarios. For example, median plasma PFOS concentrations in a highly exposed population of firefighters from Queensland, Australia (n = 149) were approximately 66 µg/L (0.13 µM), and only ∼7-fold lower than our lowest experimental dose [37].

#### 2.2.2 INS-1 832/3 cell culture

INS-1 832/3 cells (generously provided by Dr. Mathieu Ferron, Institut de recherches cliniques de Montreal), an immortalized pancreatic insulinoma beta cell line, were cultured in RPMI 1640 + L-glutamine (Corning, #MT10040CV; Tewsbury, MA, USA) supplemented with 10% heat-inactivated Fetal Bovine Serum (FBS; Sigma-Aldrich, #F1051-500ML), 50 µM 2-mercaptoethanol (Sigma-Aldrich, #M3148-100ML), and 10 mM HEPES (Thermo Fisher Scientific, #BP310-500, Waltham, MA, USA). Cells were maintained at 37°C and 5% CO_2_.

INS-1 832/3 cells (passage 7–11) were seeded into 24-well plates at a density of 1×10^6^ cells/well and allowed to adhere for 24-hrs prior to chemical exposure. At the start of chemical exposure, cells were approximately 70-75% confluent. Cells were exposed to DMSO (vehicle) or PFOS (1, 10 or 100 µM) as follows: 1) acutely for 2-hrs during a GSIS assay (**Figure 3A**; protocol described in *Section 2.3.2*.), or 2) for 48-hrs in culture media (prolonged exposure) and beta cell function was subsequently assessed using a standard GSIS assay (**Figure 3B**; protocol described in *Section 2.3.1*.).

**Figure 3.**
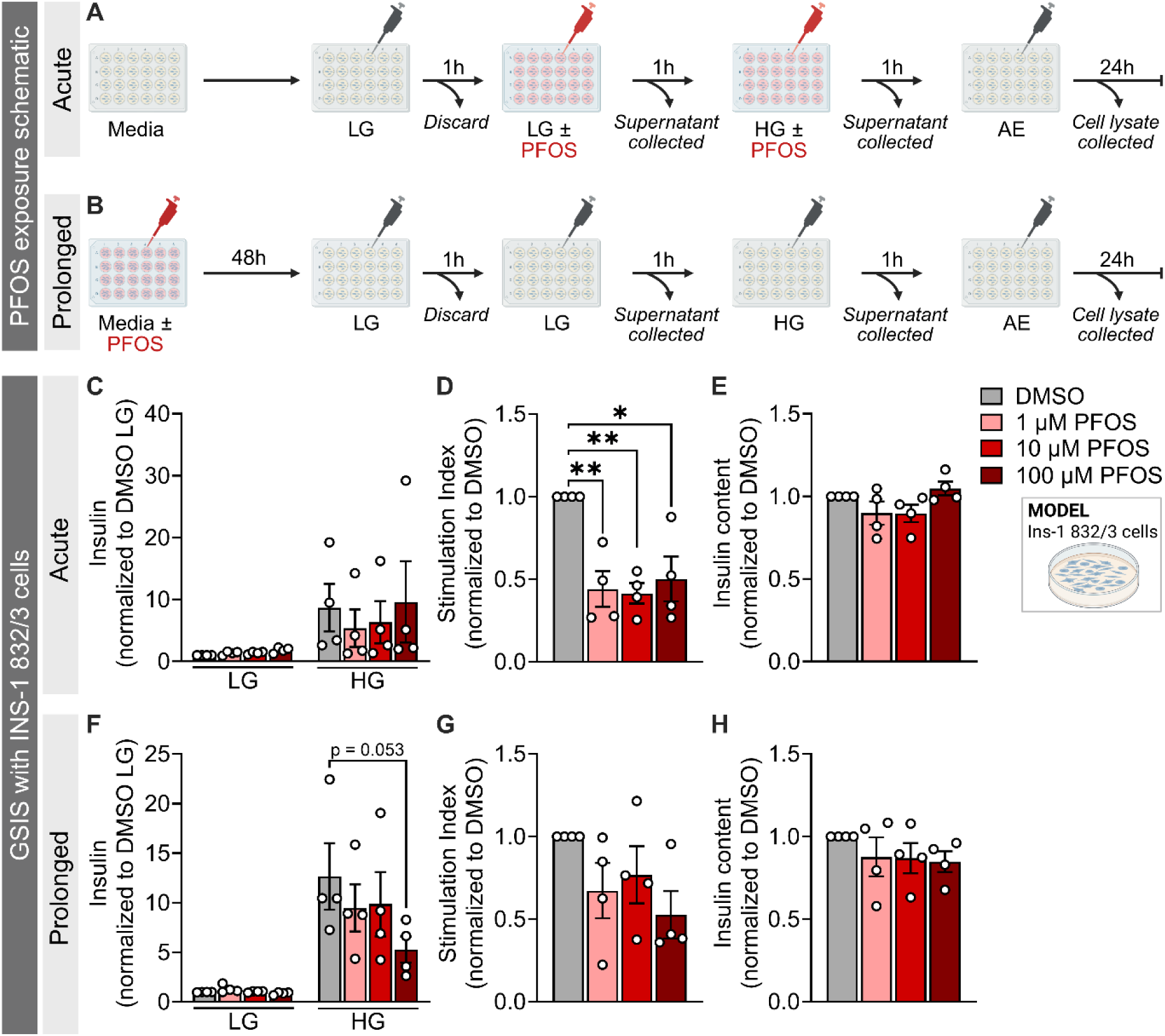
Acute and prolonged PFOS exposure suppresses insulin secretion in INS-1 832/3 cells. **(A,B)** Schematic summary outlining the method of PFOS exposure administered to immortalized INS-1 832/3 cells. Cells were exposed to vehicle (DMSO) or PFOS (1, 10, or 100 µM) either **(A)** acutely during a static glucose-stimulated insulin secretion (GSIS) assay or **(B)** for 48-hrs in culture media and cell function was subsequently assessed using a non-chemical GSIS assay. Our static GSIS protocol consisted of immersing cells in low glucose (LG; 2.8 mM) for 1-hr to pre-equilibrate the cells, followed by sequential 1-hr incubation in LG and high glucose (HG; 16.7 Mm) to assess insulin secretion. Cells were then incubation overnight in acid ethanol (AE) to assess insulin content. **(C,F)** Insulin secretion (normalized to DMSO LG), **(D,G)** stimulation index, and **(E,H)** insulin content (normalized to DMSO) following **(C-E)** acute and **(F-H)** prolonged exposure to PFOS. **(D,G)** Stimulation index was calculated as a ratio of insulin concentration under HG relative to LG. Data are presented as mean ± SEM. Individual data points represent biological replicates (n = 4–6 technical replicates per biological replicate, n = 4 biological replicates/condition). *p-value < 0.05; **p-value < 0.01. The following statistical tests were used: **(C,F)** two-way ANOVA with Tukey’s post-hoc; **(D,E,G,H)** one-way ANOVA with Dunnett’s post-hoc.

#### 2.2.3. Human islet culture

Pancreas tissue was processed for islet cell isolation (n = 4 organ donors), as described by the ADI IsletCore [38]. Isolated islets were cultured in CMRL media (Corning, #15110CV) supplemented with 0.5% BSA (30% v/v; Equitech Bio Inc, #BAL62; Kerville, TX, USA), 1% insulin-transferrin-selenium (100x; Corning, #25800CR), 2% Gibco® GlutamMAX^TM^ (100x; ThermoFisher Scientific, #35050061), and 0.5% penicillin-streptomycin (ThermoFisher Scientific, # BW09-757F). Human donor islets were shipped overnight to Carleton University (Ottawa, Ontario) in CMRL media (ThermoFisher Scientific, #11-530-037). Upon arrival, islets were transferred to low glucose Dulbecco Modified Eagle Medium (LG-DMEM; Thermo Fisher Scientific, #11885-084), supplemented with 10% FBS (Sigma-Aldrich, #F1051) and 1% penicillin-streptomycin (ThermoFisher Scientific, #15140122) and cultured overnight at 37°C and 5% CO_2_. The following morning, islets were hand-picked into treatment conditions. Islets were exposed to DMSO (vehicle) or PFOS (1, 10 or 100 µM) using the acute or prolonged treatment conditions described in *Section 2.2.2*. An experimental design schematic can be found in **Figure 4A,B**. Donor characteristics can be found in **Supplemental Table 4**.

**Figure 4.**
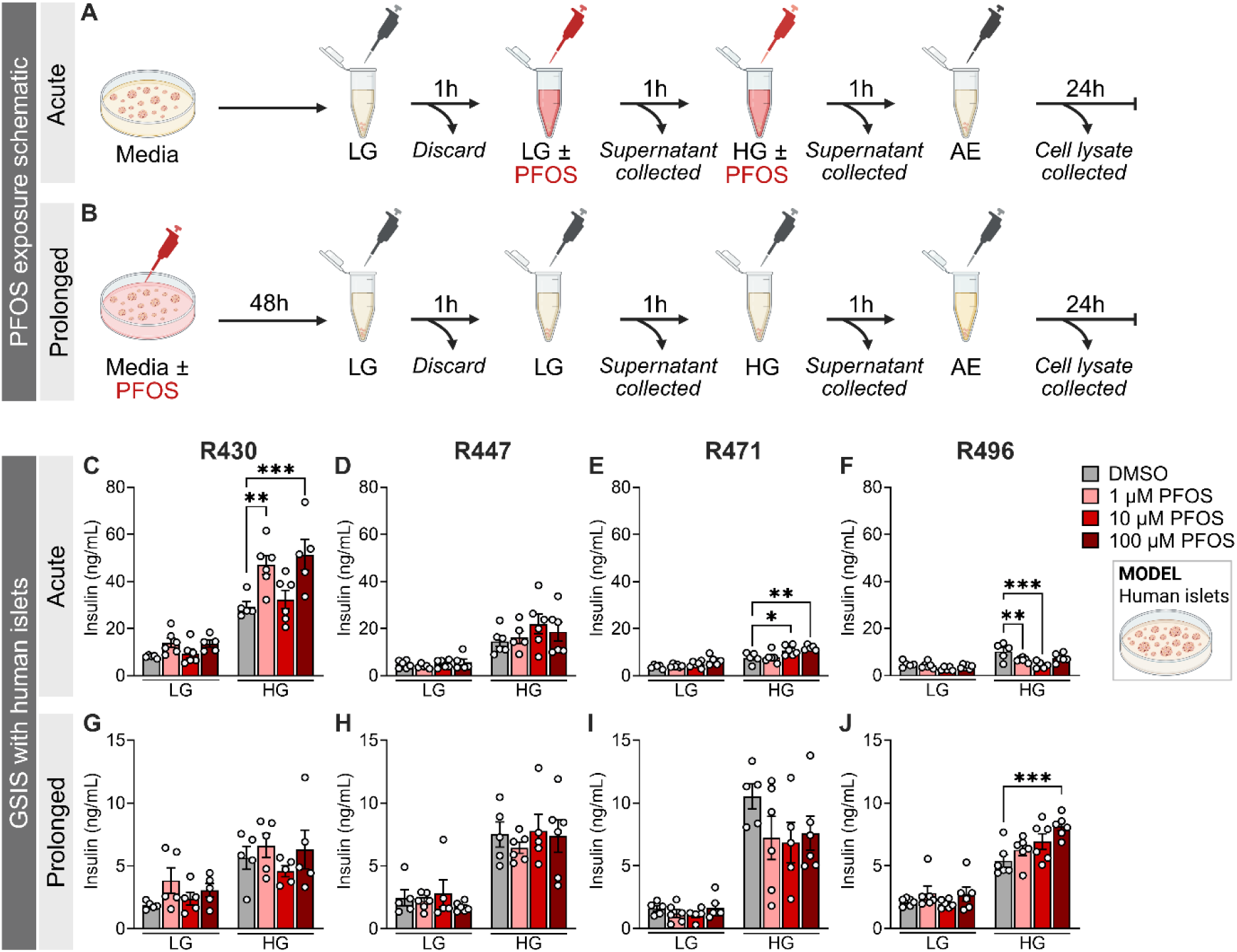
The effects of PFOS on insulin secretion in primary human islets varies between donors. **(A,B)** Schematic summary outlining the method of PFOS exposure administered to human donor islets. LG = low glucose (2.8 mM), HG = high glucose (16.7 mM). **(C-J)** Islets from 4 human organ donors were exposed to DMSO (vehicle) or PFOS (1, 10, or 100 µM) either **(C-F)** acutely during a GSIS assay or **(G-J)** for 48-hrs and islet function was subsequently assessed with a static GSIS. Data are presented as mean ± SEM. Individual data points represent technical replicates from a single donor (n = 4–6 technical replicates/donor/condition). *p-value < 0.05; **p < 0.01, ***p<0.001. The following statistical test was used: two-way ANOVA with Tukey’s post-hoc.

#### 2.2.4. Mouse islet isolation and culture

Mouse islets were isolated by pancreatic duct injection with collagenase, as previously described [39]. Information about the sex and age of the mice for each experiment is specified in *Section 2.4*. Isolated mouse islets were cultured overnight at 37°C and 5% CO_2_ in complete RPMI (Wisent, #350-000 RL; Saint-Jean-Baptiste, QC) media to allow for recovery prior to conducting experiments. The following morning, islets were exposed to DMSO (vehicle) or PFOS +/- antagonists either acutely during a dynamic perifusion assay, or for 48-hrs in culture media and islet function was subsequently assessed using a standard perifusion analysis. Detailed methodology about perifusion assessments is described in *Section 2.4*. An experimental design schematic can be found in **Figure 5A,B**.

**Figure 5.**
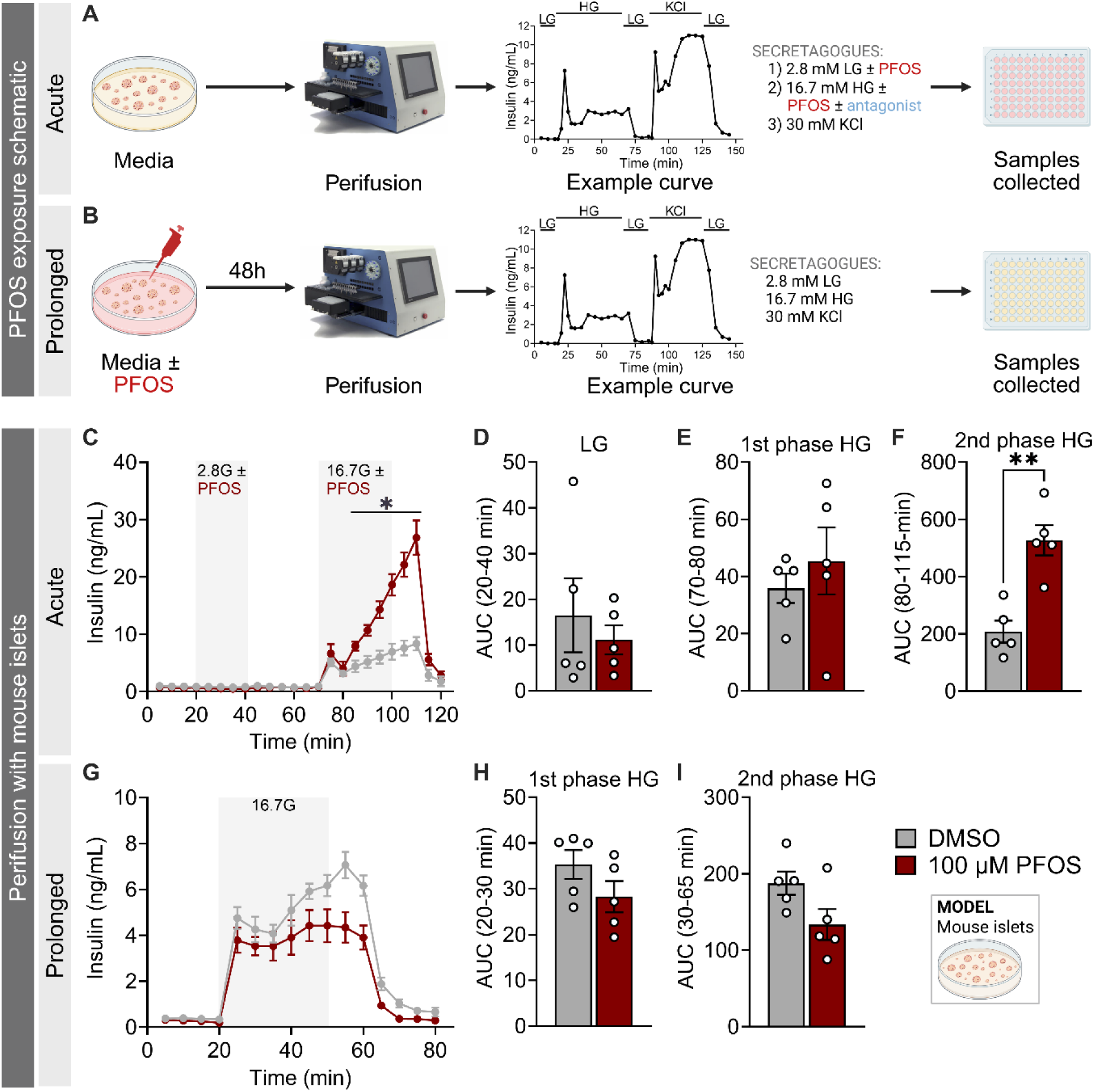
Acute high-dose PFOS exposure causes exaggerated insulin secretion in female mouse islets, whereas prolonged PFOS modestly impairs insulin secretion. **(A,B)** Schematic summary outlining the method of PFOS exposure administered to mouse islets *in vitro*. LG = low glucose, HG = high glucose (16.7 mM). Female mouse islets were exposed to DMSO (vehicle) or 100 µM PFOS **(C–F)** acutely during a GSIS assay using perifusion analysis or **(G–I)** for 48-hrs in culture media and islet function was subsequently assessed by perifusion. Data is presented as **(C,G)** dynamic insulin secretion curves and **(D–F,H,I)** area under the curve (AUC). AUCs include **(D)** the 2.8 mM low glucose (LG) stimulation, **(E,H)** 1^st^ phase GSIS, and **(F,I)** 2^nd^ phase GSIS. All data represent mean ± SEM. Individual data points on bar graphs represent biological replicates (n = 5/condition). *p-value<0.05; **p-value<0.01. The following statistical tests were used: **(C,G)** two-way repeated measures ANOVA with two-stage linear step-up procedure of Benjamini, Krieger and Yekutieli; **(D–F,H,I)** two-tailed paired t-test.

### 2.3. Static glucose-stimulated insulin secretion assay

#### 2.3.1. Static GSIS following prolonged PFOS exposure

We conducted a static GSIS assay on INS-1 832/3 cells and human donor islets exposed to either DMSO or PFOS for 48-hrs (see **Figure 3B**, **4B** for experimental designs). For adherent INS-1 832/3 cells, the GSIS assay was conducted in a 24-well culture plate (n = 5–6 technical replicates/biological replicate, n = 3 biological replicates/condition). For human islets, 25 islets per replicate were transferred to a 1.5 mL microcentrifuge tube just prior to starting the assay (n = 4–6 technical replicates/donor/condition).

Cells were washed with pre-warmed 0 mM glucose Krebs-Ringer bicarbonate buffer (KRBB; 115 mM NaCl, 5 mM KCl, 24 mM NaHCO_3_, 2.5 mM CaCl_2_, 1 mM MgCl_2_, 10 mM HEPES, 0.1% (wt/vol.) BSA, pH 7.4) to remove traces of chemical and culture media. Cells were then incubated in 2.8 mM low glucose (LG) KRBB solution for a 1-hr pre-incubation at 37°C and the supernatant was discarded. Cells were then immersed in 500 µL of 2.8 mM LG KRBB for 1-hr, followed by 500 µL of 16.7 mM high glucose (HG) KRBB for 1-hr at 37°C. The LG and HG KRBB samples were centrifuged (2000 rpm) and the supernatant stored at -30°C until analysis. Insulin content was measured by immersing cells in 500 µL of acid-ethanol solution (1.5% (vol./vol.) HCl in 70% (vol./vol.) ethanol) at 4°C overnight and then neutralizing with equal volumes of 1M Tris base (pH 7.5). Insulin concentrations were measured by ELISA (human islets: ALPCO, #80-INSHU-CH10; INS-1 832/3 cells: ALPCO, #80-INSMR-CH10; Salem, New Hampshire, USA).

#### 2.3.2. Acute PFOS exposure during a static GSIS

To assess the acute effects of PFOS on insulin secretion, we conducted a static GSIS as described in *Section 2.3.1.* but with modifications. First, cells were chemically naïve at the start of the GSIS assay. Second, after the 1-hr pre-incubation with 2.8 mM LG KRBB, we added DMSO or PFOS directly to the LG and HG KRBB solutions such that the cells were acutely exposed to PFOS during the sequential 1-hr LG and 1-hr HG incubations (cumulative exposure of 2-hrs to PFOS; see **Figure 3A**, **4A** for experimental designs).

### 2.4. Perifusion assay for dynamic insulin secretion analysis in mouse islets

We assessed dynamic insulin secretion by perifusion (Biorep Technologies, Miami Lakes, FL) in isolated mouse islets; an experimental design schematic can be found in **Figure 5A,B**. Per biological replicate, 70 islets were handpicked and loaded into Perspex microcolumns, positioned between two layers of acrylamide-based microbeads (Biorep Technologies, #PERI-BEADS-20). Islets were perifused at a flowrate of 40 µL/min or 80 µL/min depending on the experiment and the stimulation step. Throughout the perifusion process, the islets and perifusion solutions were maintained at 37°C in a temperature-controlled chamber while the collection plate was maintained at 4°C using the tray cooling system. Samples were stored at -80°C until further analysis. Insulin concentrations were measured for all experiments using an ELISA (ALPCO, #80-INSMR-CH10). Glucagon concentrations were measured in DMSO and PFOS-exposed samples from *Section 2.4.4. Acute PFOS exposure +/- Ex(9-39)* using the Promega Lumit® Glucagon Immunoassay (Promega, #W8020; Madison, Wisconsin, USA).

#### 2.4.1. Prolonged and acute PFOS exposure in female islets

C57Bl/6J female mice were bred at Carleton University (16–18 weeks old; n = 6 biological reps/condition). Islets were exposed to 0.1% DMSO or 100 µM PFOS for 48-hrs in culture media (**Figure 5G-I**) or acutely during perifusion analysis (**Figure 5C-F**).

Islets were washed 1X with PBS(+/+) to remove traces of chemical and/or media prior to running perifusion. Islets were perifused for 40 min with 2.8 mM LG KRBB to equilibrate the islets (samples discarded). Following 48-hr PFOS exposure, islets were perifused with the following protocol: (i) LG KRBB for 15 min, (ii) HG KRBB for 35 min, and (iii) LG KRBB for 20 min. For acute PFOS exposure during the perifusion assay, islets were perifused with the following protocol: (i) LG (no chemical) for 15 min, (ii) LG KRBB with either DMSO or PFOS for 25 min, (iii) LG (no chemical) for 25 min, (iv) HG KRBB containing either DMSO or PFOS for 35 min, and (v) LG KRBB (no chemical) for 20 min.

#### 2.4.2. Acute PFOS dose-response in male and female islets

C57Bl/6J female and male mice were purchased from Charles River or bred at Carleton University (9–12 weeks old; 3–6 biological reps/sex/condition). Islets were acutely exposed to 0.1% DMSO or PFOS (1, 10, 100 µM) during the perifusion analysis (**Figure 6**; see **Figure 5A** for experimental design schematic). Female and male isIets were perifused for 40 min with 2.8 mM LG KRBB to equilibrate the islets (samples discarded). Female islets were then perifused with the following protocol: (i) LG KRBB for 15 min, (ii) HG KRBB containing either DMSO or PFOS for 65 min, (iii) LG KRBB for 20 min, (iv) KCl KRBB for 35 min, and (v) LG KRBB for 15 min. Male islets were perifused with the following protocol: (i) LG KRBB for 15 min, (ii) HG KRBB containing either DMSO or PFOS for 45 min, (iii) LG KRBB for 25 min, (iv) KCl KRBB for 35 min, and (v) LG KRBB for 25 min.

**Figure 6.**
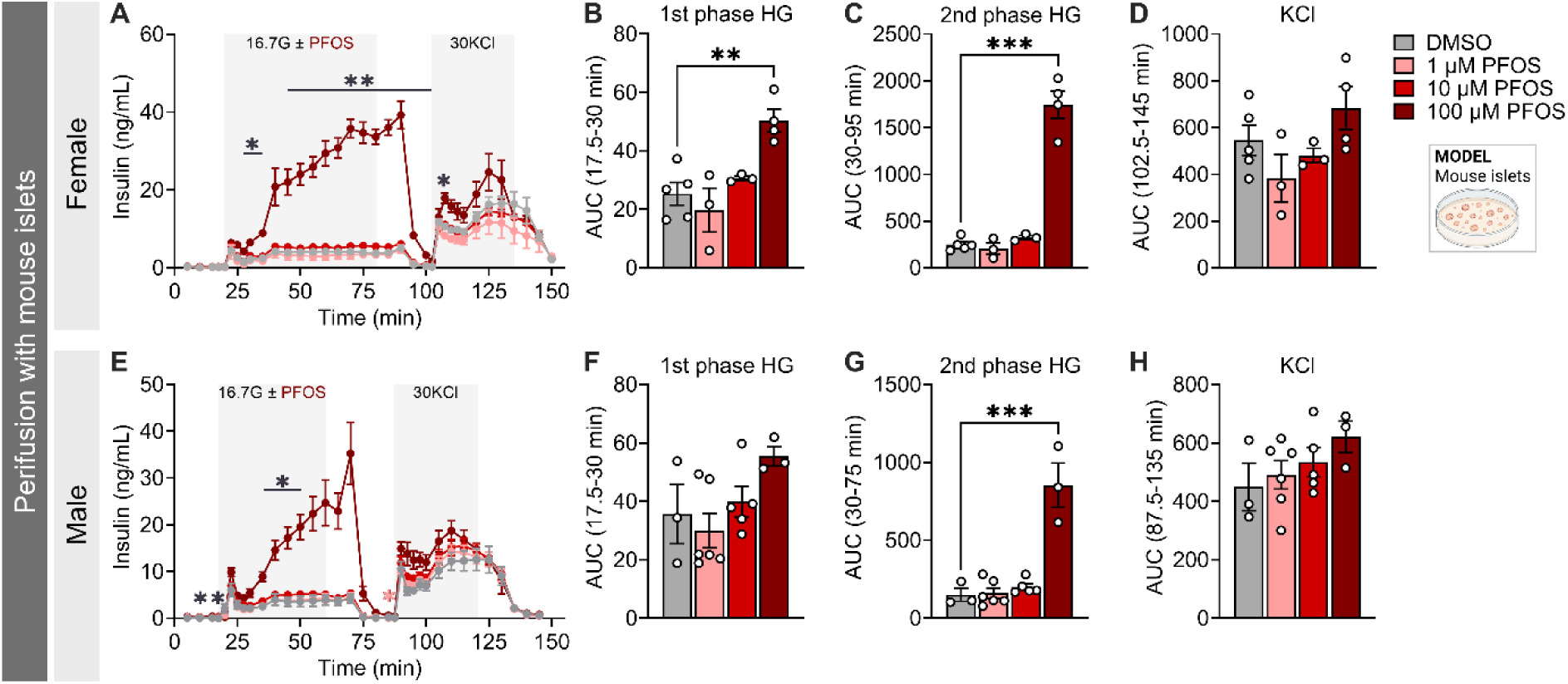
PFOS acutely stimulates insulin secretion under high glucose conditions in female and male mouse islets. **(A-D)** Female and **(E-H)** male mouse islets were acutely exposed to DMSO (vehicle) or PFOS (1, 10, 100 µM) during a GSIS assay using perifusion analysis (see Figure 5A**,B** for experimental design schematic). High glucose (HG) = 16.7 mM (16.7G), KCl = 30 mM. **(A,E)** Dynamic insulin secretion curves in response to LG (2.8 mM), HG (16.7 mM ± DMSO or 1-100 µM PFOS), and KCl (30 mM). Area under the curve for **(B,F)** 1^st^ phase GSIS **(C,G)** 2^nd^ phase GSIS, and **(D,H)** the KCl stimulation step. All data represent mean ± SEM. Individual datapoints represent biological replicates (n = 3–6 mice/sex/condition). *p<0.05, **p<0.01. The following statistical tests were used: **(A,E)** two-way repeated measures ANOVA with Dunnett’s post-hoc; **(B–D,F–H)** one-way ANOVA with Dunnett’s post-hoc.

#### 2.4.3 Acute PFOS exposure +/- GW1100

The GPR40 antagonist GW1100 (Millipore Sigma, #371830; Oakville, Ontario, Canada) was solubilized in DMSO to a final stock concentration of 50 mM. The solution was kept at -20°C for long-term storage. Islets were acutely exposed to 0.1% DMSO or 100 µM PFOS with or without 30 µM GW1100 (**Figure 7A-B**; see **Figure 5A** for experimental design schematic). Female C57BL/6J mice were purchased from Jackson Laboratories (6–10 weeks old; n = 3 biological reps/condition). Islets were perifused with LG KRBB +/- GW1100 for 40 min to equilibrate the islets (samples discarded). Islets with then perifused with the following protocol: (i) LG KRBB +/- GW1100 for 15 min, (ii) HG KRBB containing either DMSO, PFOS, GW1100, or GW1100 + PFOS for 65 min, (iii) LG KRBB (no GW1100) for 15 min, (iv) KCl KRBB for 35 min, and (v) LG KRBB (no GW1100) for 15 min.

**Figure 7.**
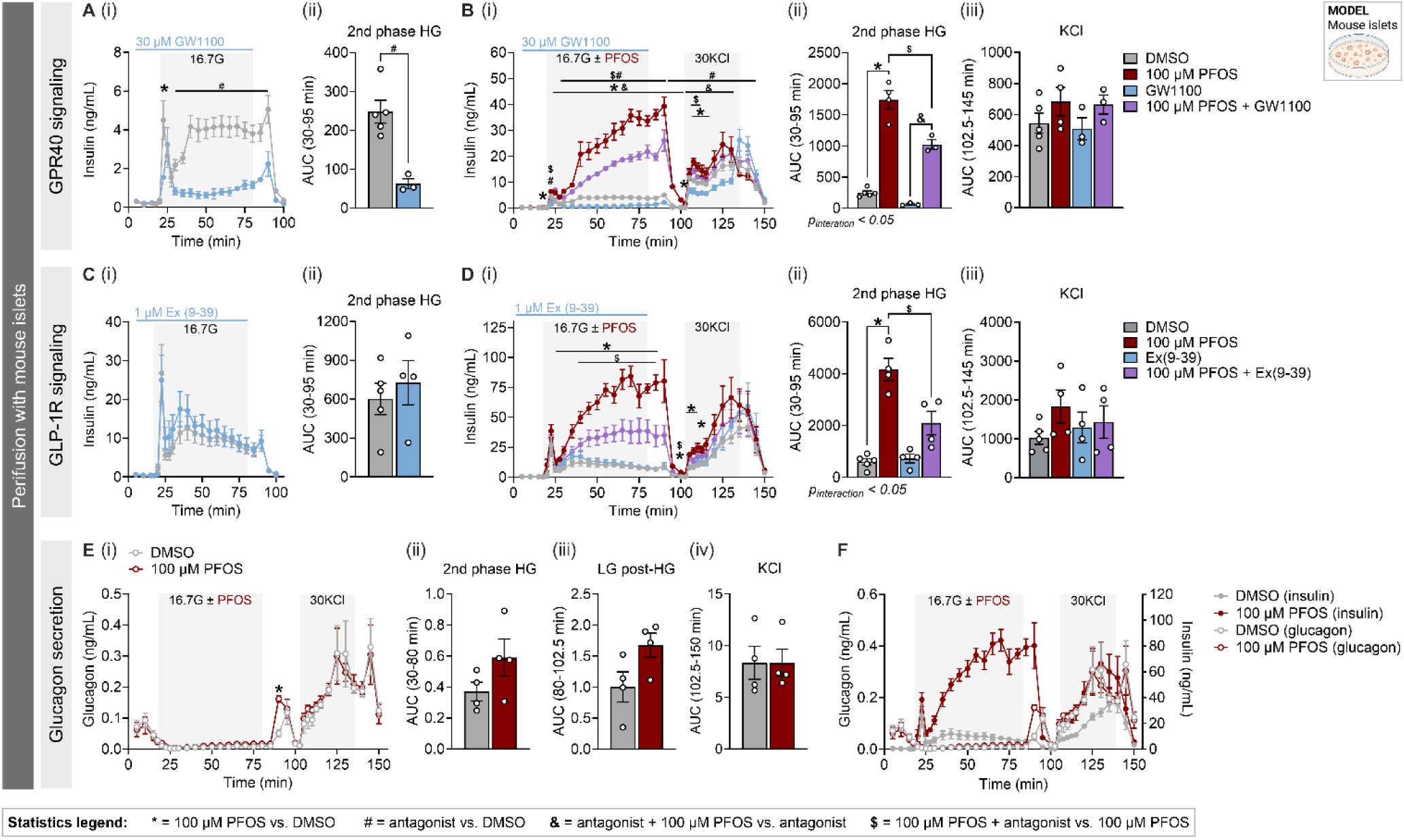
PFOS-induced hyperinsulinemia is partially mediated by GLP-1R, but not GPR40, in female mouse islets. **(A-D)** Female mouse islets were acutely exposed to DMSO (vehicle) or 100 µM PFOS with or without **(A,B)** a GPR40 antagonist (GW1100), and **(C,D)** a GLP-1R antagonist (Ex9-39) during a perifusion analysis (see Figure 5A**,B** for experimental design schematic). High glucose (HG) = 16.7 mM (16.7G), KCl = 30 mM. **(A,C)** Perifusion data comparing insulin secretion following exposure to DMSO ± antagonist, presented as (i) dynamic secretion curves and (ii) area under the curve (AUC) of 2^nd^ phase GSIS. **(B,D)** Perifusion data comparing insulin secretion following exposure to **(B)** DMSO, 100 µM PFOS only, 30 µM GW1100 only, or 100 µM PFOS + 30 µM GW1100, and **(D)** DMSO, 100 µM PFOS only, 1 µM Ex(9-39) only, or 100 µM PFOS + 1 µM Ex(9-39). Data is presented as (i) dynamic secretion curves, (ii) AUC for 2nd phase GSIS, and (iii) AUC for the KCl stimulation. **(E)** Glucagon secretion following exposure to DMSO or 100 µM PFOS, presented as (i) dynamic secretion curves, (ii) AUC of 2^nd^ phase GSIS, (iii) AUC for LG incubator that follows the HG stimulation, and (iv) AUC for the KCl stimulation. **(F)** Overlay of the perifusion curves showing insulin secretion and glucagon secretion (data extracted from panels **D** and **E**). A p-value < 0.05 was considered significant and was denoted by the following symbols: * = 100 µM PFOS vs. DMSO; # = antagonist vs. DMSO; & = antagonist + 100 µM PFOS vs. antagonist; $ = 100 µM PFOS + antagonist vs. 100 µM PFOS. Individual datapoints represent biological replicates (n = 3–5 mice/condition). The following statistical tests were used: **(Ai,Ci)** two-way repeated measures ANOVA with False Discovery Rate post-hoc correction (two-stage step-up method of Benjamini, Krieger and Yekutieli); **(Aii,Cii)** unpaired two-tailed t-test; **(Bi,Di)** three-way repeated measures ANOVA with False Discovery Rate post-hoc correction (two-stage step-up method of Benjamini, Krieger and Yekutieli), **(Bii,Biii,Cii,Ciii)** two-way ANOVA with Tukey’s post-hoc test; **(Ei)** two-way repeated measures ANOVA with False Discovery Rate post-hoc correction (two-stage step-up method of Benjamini, Krieger and Yekutieli); **(Eii,Eiii,Eiv)** two-tailed paired t-test

#### 2.4.4. Acute PFOS exposure +/- Ex(9-39)

The GLP-1 receptor antagonist (Ex)9-39 amide (BACHEM Holding, #4017799; Bubendorf, Switzerland) was solubilized in DMSO, to a final stock concentration of 0.3 mM. The solution was kept at -20°C for long-term storage. Islets were acutely exposed to 0.1% DMSO or 100 µM PFOS with or without 1 µM (Ex)9-39 (**Figure 7C-D**; see **Figure 5A** for experimental design schematic).

Female C57Bl/6J mice were bred at Carleton University (21 weeks old; n = 3 biological reps/condition). Islets were perifused with LG KRBB for 40 min to equilibrate the islets (samples discarded). Islets with then perifused with the following protocol: (i) LG KRBB +/- Ex(9-39) for 15 min, (ii) HG KRBB containing DMSO, PFOS, Ex(9-39), or Ex(9-39) + PFOS for 65 min, (iii) LG KRBB (no Ex(9-39)) for 15 min, (iv) KCl KRBB for 35 min, and (v) LG KRBB (Ex(9-39)) for 15 min.

### 2.5. Acute effects of in vivo PFOS exposure with or without a glucose bolus

All mice received ad libitum access to standard rodent chow (Harlan Laboratories, Teklad Diet #2014; Madison, WI) and were maintained on a 12-hour light/dark cycle. All experiments were approved by the Carleton University Animal Care Committee and carried out in accordance with the Canadian Council on Animal Care guidelines. Prior to starting the study, animals were randomly assigned to treatment groups and matched for body weight and blood glucose to ensure that these variables were consistent between experimental groups.

Female C57Bl/6J mice (6–27 weeks-old; n = 16/condition) were bred at Carleton University. All mice underwent a 6-hr morning fast prior to metabolic assessment. A subset of mice received an injection of DMSO (vehicle) or 10 mg/kg PFOS via intraperitoneal injection (study schematic in **Figure 8A**). Another subset received a bolus of glucose (2 g/kg) containing DMSO (vehicle) or 10 mg/kg PFOS via intraperitoneal injection (study schematic in **Figure 8A**). Blood glucose was measured at t = 0, 2, 7, 15, 30, and 60 min using a handheld glucometer (OneTouch Verio, #023-194). Plasma was collected at t = 0, 2, 7, and 15 min using heparinized microhematocrit tubes and plasma insulin levels were measured by ELISA (ALPCO; 80-INSMU-CH10). Time 0 indicates the blood sample collected prior to administration of glucose. All assessments were performed in conscious, restrained mice, with blood samples collected via the saphenous vein.

**Figure 8.**
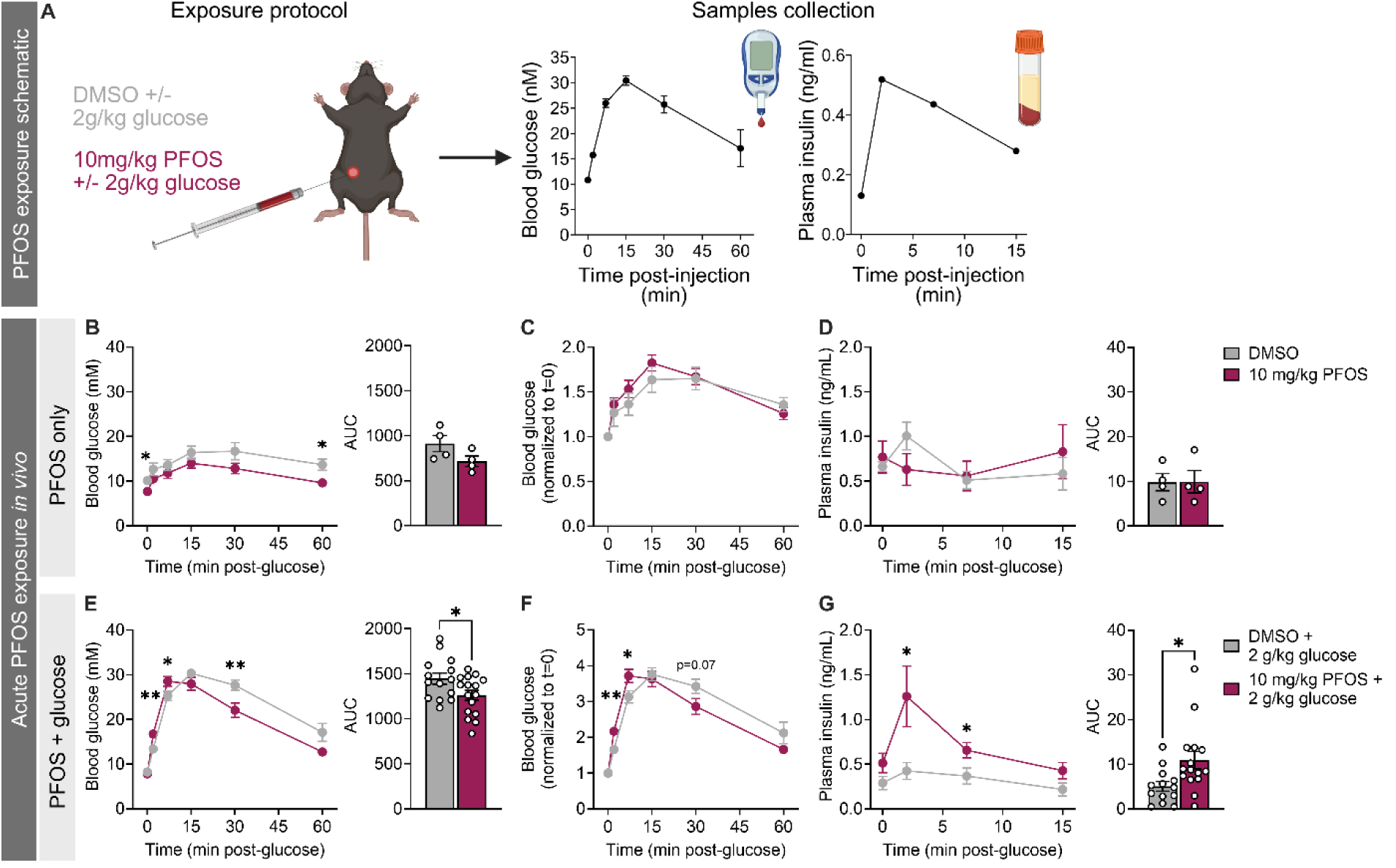
Acute PFOS exposure during a glucose tolerance test *in vivo* stimulates insulin secretion in female mice. **(A)** Schematic summary timeline. Female mice were administered either DMSO (vehicle) or 10 mg/kg PFOS via intraperitoneal injection in the absence of glucose (n = 4 mice/condition) or concurrently with 2 g/kg glucose during a glucose tolerance test (n = 13–16 mice/condition). **(B,C,E,F)** Blood glucose, and **(D,G)** plasma insulin. Data are presented as a line graph and area under the curve. Blood glucose data is presented as **(B,E)** raw data, and **(C,F)** data normalized to baseline blood glucose (at t = 0). All data are presented as mean ± SEM. Individual data points on bar graphs represent different mice. The following statistical tests were used: **(B–G** line graph) two-way repeated measures ANOVA with Šídák’s multiple comparisons test; **(B–G AUC)** two-tailed unpaired t-test.

Although supraphysiological, the 10 mg/kg dose was selected to align with the dose used in the Qin et al. study in which they reported that PFOS exposure significantly increased insulin secretion 1-hr post-gavage in male mice [21]. This dose should generate mouse plasma PFOS concentrations of ∼2800 µM (assuming an average body weight of 25 g and an average injection volume of 0.18 mL).

### 2.6. Quantification and Statistical Analysis

All statistics were performed using GraphPad Prism 9.1.1 (GraphPad Software Inc., La Jolla, CA). For static GSIS and perifusion data, a two-way ANOVA was used with the Tukey’s post-hoc test. For our antagonist perifusion data, a three-way ANOVA was used with the false discovery rate (FDR) post-hoc test. For GTT data, a two-way repeated measures ANOVA was used with Tukey’s post-hoc test. AUC bar plots were analysed with a one-way or two-way ANOVA depending on the experimental design. Additional information on statistical tests can be found in figure legends. Significance was determined as p<0.05. Outliers were identified using the extreme studentized deviate method.

## 3. RESULTS

### 3.1. PFOS is detected in human and mouse pancreas, and at higher concentrations than other legacy PFAS analytes

We measured PFOA, PFOS and PFHxS concentrations in human pancreas and plasma from 88 and 79 donors, respectively. PFOS was detected in 99% of pancreas samples (87/88 donors), compared to 70.4% for PFHxS and only 16% for PFOA; all three legacy PFAS were detected in 100% of plasma samples (**Figure 1A**). Interestingly, PFOS was detected at higher concentrations in both pancreas and plasma compared to PFOA or PFHxS (**Figure 1B,C**). Although most donors had higher levels of PFOS in plasma compared to pancreas, we identified a subset of donors (8 of 79) that had higher PFOS concentrations in pancreas compared to plasma (**Figure 1D**). These data suggest that PFOS accumulates more readily in pancreas compared to other legacy PFAS analytes, and in some donors, at higher concentrations than in plasma.

To confirm if PFOS and PFOA have different affinities for accumulating in pancreas tissue, we modelled human PFAS exposure in mice. Mice received daily oral exposure to 0.5% Tween-20 (vehicle) or a supraphysiological dose of either 4.5 mg/kg PFOS or 4.5 mg/kg PFOA for 28 days (**Figure 2A**). Interestingly, PFOS-exposed mice had a median PFOS concentration of 28,000 ng/g in pancreas, whereas PFOA-exposed mice only had 15,000 ng/g of PFOA in pancreas (**Figure 2B**). Therefore, despite having been exposed to the same concentration of either PFOS or PFOA for the same duration of time, PFOS accumulated in pancreas at concentrations twice as high as PFOA.

We next exposed female mice to environmentally relevant doses of PFOS (0.6, 60 µg/L) and a supraphysiological dose of PFOS (6000 µg/L) via drinking water for 21 days (**Figure 2C**) to determine if these doses would result in biologically meaningful PFOS accumulation in pancreas. We detected measurable levels of PFOS in the pancreas after exposing mice to a dose as low as 0.6 µg/L (range: 0.2– 5.1 ng/g) and 60 µg/L (range: 25.2–28.5 ng/g; **Figure 2D**), which generally aligned with the concentrations of PFOS detected in human pancreas from our cohort of 88 human donors (range: 0.083–5.5 ng/g; **Figure 1C**). Collectively, these data confirm that the pancreas is a site for PFAS accumulation and that PFOS levels are of particular concern, emphasizing the importance of understanding the effects of PFOS on beta cell function.

### 3.2. Acute and prolonged PFOS exposure suppresses insulin secretion in INS-1 832/3 cells

We first tested the impact of PFOS on beta cell function using immortalized rat INS-1 832/3 cells as a readily available controlled model system, and to build on previous studies in immortalized beta-TC-6 cells [20–22]. We exposed INS-1 832/3 cells to PFOS for either 2-hrs during a static GSIS (acute exposure; **Figure 3A**) or for 48-hrs in culture media (prolonged exposure) prior to conducting a static GSIS assay (**Figure 3B**). Acute PFOS exposure had no effect on insulin secretion under LG or HG conditions (**Figure 3C**), or on total insulin content (**Figure 3E**), but significantly suppressed the stimulation index (HG:LG ratio) at all doses compared to DMSO (**Figure 3D**). Prolonged PFOS exposure caused a trending decrease in insulin secretion under HG conditions (p = 0.053, **Figure 3F**), but only at the highest PFOS dose (100 μM). Prolonged PFOS exposure had no effect on stimulation index (**Figure 3G**) or insulin content (**Figure 3H**). Collectively, these data indicate that PFOS can directly impair beta cell function in INS-1 832/3 cells, with acute exposure having more robust effects than prolonged exposure.

### 3.3. The effects of PFOS on insulin secretion in primary human islets varies between donors

Given the differences between immortalized beta cells and intact primary islets [40], we next assessed the effect of PFOS exposure on primary human islets isolated from deceased organ donors as a more physiologically relevant model system. For each donor, we exposed a subset of islets to PFOS either acutely during a static GSIS (**Figure 4A**) or for 48-hrs in culture media (**Figure 4B**).

We observed substantial biological variability in the response of donors to both acute and prolonged PFOS exposure. Following acute PFOS exposure, donors R430 and R471 exhibited a statistically significant increase in insulin secretion under HG conditions (**Figure 4C,E**), donor R496 showed suppressed insulin secretion under HG conditions (**Figure 4F**), and donor R447 displayed no change in insulin secretion under HG conditions (**Figure 4D**). Acute PFOS exposure did not impact insulin secretion under LG conditions (**Figure 4C-F**), stimulation index (**Supplemental Figure 1A,E,I,M**), or insulin content (**Supplemental Figure 1B,F,J,N**) in any of the donors.

Prolonged PFOS exposure had no impact on insulin secretion under either LG or HG conditions (**Figure 4G-J**), except for R496, which displayed a modest increase in GSIS following 100 μM PFOS exposure (**Figure 4J**). There were no changes in stimulation index (**Supplemental Figure 1C,G,K,O**) or total insulin content (**Supplemental Figure 1D,H,L,P**) following prolonged PFOS exposure.

### 3.4. Acute high-dose PFOS exposure causes exaggerated insulin secretion in female mouse islets, whereas prolonged PFOS modestly impairs insulin secretion

Due to the biological variability between human donors, we were limited in our ability to draw clear conclusions about the acute and prolonged effects of PFOS on islet function. As such, we next tested the effects of PFOS in primary mouse islets using more detailed perifusion analysis (**Figure 5A,B**).

Acute PFOS exposure had no effect on insulin secretion under LG conditions (**Figure 5C,D**). Interestingly, under HG conditions, acute PFOS exposure had no effect on first-phase insulin secretion (**Figure 5C,E**) but caused a dramatic 3.4-fold increase in second-phase insulin secretion relative to DMSO control (**Figure 5C,F**). In contrast, prolonged PFOS exposure caused a trending, but non-significant, decrease in both first-phase (**Figure 5G,H**) and second-phase (**Figure 5G,I**) secretion under HG conditions. Collectively, these data demonstrate that PFOS acutely amplifies the insulin response to HG but not LG.

### 3.5. PFOS acutely stimulates insulin secretion under high glucose conditions in female and male mouse islets

Next, we expanded on our findings in female mouse islets to determine whether the acute effects of PFOS on the insulin response to HG were reproducible, dose-dependent, and/or sex-specific. We also extended the duration of the HG stimulation to determine whether the stimulatory effects of PFOS on insulin secretion would eventually plateau and/or lead to insulin depletion and reduced insulin secretion. We added a KCl stimulation (no chemical) post-HG to determine if there were lingering effects on insulin secretion capacity after the removal of PFOS. In summary, islets isolated from both female and male mice were exposed to various doses of PFOS (1, 10, 100 μM) during the HG stimulation, followed by a KCl stimulation in the absence of chemical (**Figure 6**).

Importantly, we were able to reproduce our findings showing that 100 μM PFOS caused an ∼3-fold increase in insulin secretion under HG conditions in female mouse islets, with the most robust effect occurring during second-phase insulin secretion (**Figure 6A–C**). There was no overall effect of 100 μM PFOS on insulin secretion capacity during the KCl stimulation (**Figure 6A,D**). Interestingly, 1 and 10 μM PFOS did not impact glucose- or KCl-stimulated insulin secretion in female islets (**Figure 6A–D**). In male islets, 100 µM PFOS also caused a robust increase in insulin secretion under HG conditions (**Figure 6E–G**), with no effect on KCl-stimulated insulin secretion (**Figure 6E,H**). These data confirm that acute exposure to 100 µM PFOS profoundly hyperstimulates second-phase insulin secretion under HG conditions, irrespective of sex.

### 3.6. PFOS-induced hyperinsulinemia is partially mediated by GLP-1R but not GPR40, in female mouse islets

Due to structural similarities between long-chain fatty acids—the natural ligands for GPR40—and PFOS, previous studies hypothesized that PFOS binds to GPR40 [20,21], which could explain the amplification of second-phase GSIS we observed in our perifusion (**Figure 5,6**). To evaluate this, we replicated our perifusion analysis in female mouse islets using 100 µM PFOS alongside a GPR40 antagonist, GW1100, to generate the following experimental groups: DMSO (vehicle), 100 µM PFOS only, 30 µM GW1100 only, and 100 µM PFOS + 30 µM GW1100 (**Figure 7A,B**). We again show that 100 µM PFOS robustly stimulates second-phase insulin secretion compared to DMSO in female mouse islets (**Figure 7Bi,ii**). We observed a diminished effect of PFOS on insulin secretion when islets were co-exposed to GW1100 (i.e. PFOS + GW1100 vs PFOS alone; **Figure 7Bi,ii**), however this decrease was proportional to the suppressed insulin secretion caused by GW1100 alone (i.e. GW1100 vs DMSO; **Figure 7Ai,ii**). In fact, the magnitude of insulin hypersecretion observed under the PFOS condition relative to the DMSO control (10-fold) was smaller than the magnitude of hypersecretion under the PFOS + GW1100 condition relative to the GW1100 control (20-fold) (**Figure 7Bi,ii**). There was no effect of PFOS +/- GW1100 on KCl-induced insulin secretion (**Figure 7Bi,iii**). Collectively, these data suggest that GPR40 is not driving the acute effects of PFOS on insulin secretion in female mouse islets.

Given the discrepancy in the acute effects of PFOS to suppress GSIS in INS-1 823/3 cells (**Figure 3C–E**) but amplify GSIS in mouse islets (**Figure 5**–**6**) and the finding that PFOS only alters second-phase insulin secretion in mouse islets (**Figure 5**–**6**), we next explored the possibility that PFOS alters insulin secretion via paracrine signaling. We focused on GLP-1R since this is a surface receptor on the beta cell that is known to potentiate GSIS when bound by glucagon or GLP-1 secreted from alpha cells [41,42]. To assess this, we replicated our perifusion analysis in female islets using 100 µM PFOS alongside a GLP-1R antagonist, Ex(9-39), to generate the following experimental groups: DMSO (vehicle), 100 µM PFOS only, 1 µM Ex(9-39) only, and 100 µM PFOS + 1 µM Ex(9-39) (**Figure 7C,D**). Once again, 100 µM PFOS acutely hyper-stimulated second-phase insulin secretion (**Figure 7Di,ii**). Interestingly, this phenotype was diminished by ∼2-fold when PFOS was co-administered with Ex(9-39) (i.e. PFOS only vs PFOS + Ex(9-39)), but not completely abolished (**Figure 7Di,ii**). Importantly, there was no significant effect of Ex(9-39) alone on glucose-induced insulin secretion (**Figure 7Ci,ii**) and no change in KCl-induced insulin secretion in any of the groups (**Figure 7Diii**). Based on this, we speculated that PFOS may acutely induce glucagon secretion, which would subsequently stimulate GLP-1R and amplify GSIS. To test this hypothesis, we measured glucagon secretion in islets acutely exposed to DMSO and 100 µM PFOS during the perifusion. In contrast to insulin secretion, there was no effect of PFOS on glucagon secretion under HG conditions (**Figure 7Ei,ii**). However, we observed a modest hypersecretion of glucagon immediately following the HG stimulation (t = 90 min) in the PFOS exposure group compared to DMSO (**Figure 7Ei,iiI**), which coincided with the decrease in insulin secretion post HG (**Figure 7F**). Collectively, these data suggest that the effects of PFOS on GSIS are partially mediated by GLP-1R, but not via changes in glucagon secretion.

### 3.7. Acute PFOS exposure during a glucose tolerance test *in vivo* stimulates insulin secretion in female mice

Lastly, we assessed whether the acute effects of PFOS on GSIS observed *in vitro* are reproducible in an *in vivo* mouse model. Female mice were administered DMSO or 10 mg/kg PFOS either with or without a 2 g/kg glucose bolus intraperitoneally, and blood glucose and plasma insulin levels were assessed for up to 60 min post injection (**Figure 8A**). When PFOS was administered alone, PFOS-exposed mice had slightly lower blood glucose levels at 60-min following injection (**Figure 8B**), but this is likely explained by the modest decrease in baseline fasting glucose (t = 0) in this group relative to DMSO-exposed mice (**Figure 8B**); there was no effect of PFOS on glucose tolerance when blood glucose was normalized to baseline (**Figure 8C**). Interestingly, mice simultaneously exposed to PFOS alongside a glucose bolus showed modest hyperglycemia at 2- and 7-min post injection (**Figure 8E,F**), but then transitioned to hypoglycemia at 30-min compared to DMSO control, leading to an overall decrease in blood glucose during the GTT (**Figure 8E – AUC**).

Importantly, mice exposed to PFOS with glucose showed a significant increase in plasma insulin levels at 2- and 7-min post-injection compared to mice exposed to DMSO with glucose (**Figure 8G**), whereas PFOS exposure alone (no glucose) had no effect on plasma insulin (**Figure 8D**). Collectively, these data align with our *in vitro* perifusion data showing that PFOS stimulates insulin secretion but only with simultaneous co-exposure to high glucose.

## 4. DISCUSSION

In the present study, we report that legacy PFAS chemicals are consistently detected in human pancreas tissue from a cohort of 88 organ donors; PFOS was detected in 99% of pancreas samples. We also report that PFOS concentrations were higher in pancreas compared to plasma in ∼10% of the human donors from our study. This has important implications for pancreatic health and emphasizes the relevance of understanding how PFAS impact metabolic homeostasis. We provide compelling evidence that PFOS exposure, at least at high doses, dysregulates insulin secretion in various model systems, although effects varied based on the model and duration of exposure. Prolonged PFOS exposure decreased GSIS in INS-1 832/3 cells and mouse islets, but not in human islets. In contrast, high-dose PFOS acutely stimulated second-phase GSIS in female and male mouse islets, an effect that was diminished in the presence of a GLP-1R antagonist. PFOS also acutely hyperstimulated GSIS in 2 of 4 human islet donors but not in INS-1 832/3 cells. Importantly, acute PFOS exposure *in vivo* robustly increased glucose-stimulated plasma insulin levels in female mice but only when co-administered with glucose. Collectively, these data indicate that PFOS can acutely dysregulate insulin secretion in mouse islets, likely due in part to altering paracrine signalling. Our study also demonstrates the importance of model selection when assessing toxicological effects of PFAS on islet function.

We found that PFOS is present at higher concentrations than both PFOA and PFHxS in human pancreas and plasma. Specifically, PFOS concentrations were 2.3-fold and 2.5-fold higher than PFOA in plasma and pancreas, respectively. Our data aligns with a study by Maestri et al. that measured PFOA and PFOS in human tissues, pooled from 7 donors [19]. They reported that PFOS concentrations were 1.5- and 2.5-fold higher than PFOA in both plasma and pancreas, respectively [19]. Maestri et al. also reported that PFOS concentrations were 4.4-fold higher than PFOA in liver, 1.2-fold higher in adipose, 1.8-fold higher in kidney, and 1.4-fold higher in thyroid [19]. These data suggest preferential bioaccumulation of PFOS in metabolically relevant tissues compared to other legacy PFAS. We confirmed the findings in pancreas using a controlled mouse model where the same dose and duration of PFOS or PFOA was administered over a 28-day period, yet PFOS accumulated at ∼2-fold higher concentrations in pancreas compared to PFOA. We also demonstrated that PFOS accumulates in mouse pancreas tissue following exposure to environmentally relevant doses of PFOS in drinking water (0.6 and 60 µg/L). Collectively, these findings emphasize that PFOS readily bioaccumulates in pancreas, which may have a direct impact on pancreas function. Future studies should assess a broader range of PFAS—including precursor and replacement PFAS—for potential accumulation in pancreas tissue.

This is the first study to evaluate the effects of PFOS exposure on primary mouse islets. Previous findings in immortalized beta-TC-6 cells showed insulin hypersecretion after a 1-hr exposure to 50-200 mM PFOS under 1.4 mM LG or 16.7 mM HG conditions [20,21]. Surprisingly, acute PFOS exposure had no effect on insulin secretion under LG or HG conditions in immortalized INS-1 832/3 cells in our study but rather suppressed insulin stimulation index; this may point to inherent differences between INS-1 832/3 and beta-TC-6 cells. Additionally, stimulation index was not assessed in previous studies since sequential LG and HG conditions were not evaluated. However, aligning with previous findings [20,21], we repeatedly show that acute PFOS causes an increase in GSIS in mouse islets during a perifusion analysis, but with no change in basal insulin secretion under LG condition. Importantly, these findings translated to an *in vivo* model, whereby female mice exposed to 10 mg/kg PFOS showed significantly elevated plasma insulin levels as early as 2-min post-injection, but only when PFOS was co-administered with glucose. Collectively, we provide convincing evidence that acute PFOS exposure causes hypersecretion of insulin under HG but not LG conditions in mouse islets.

Although PFOS only amplified GSIS at supraphysiological doses, these findings are concerning since they suggest that dietary intake of PFAS-contaminated food/water could exacerbate the insulin response to a postprandial glucose spike. Repeated or continuous hypersecretion of insulin, even modest, could cause beta cell stress and exhaustion, ultimately increasing the risk of developing hyperglycemia, insulin resistance, and type 2 diabetes [3,43,44]. In fact, we show that prolonged PFOS exposure for 48-hrs *in vitro* ultimately supressed insulin secretion in both INS-1 832/3 cells and mouse islets; these data align with past studies showing suppressed GSIS following 24-hr or 48-hr exposure to 100 pM – 400 µM PFOS in immortalized beta-TC-6 cells and MIN6 cells, respectively [22,25]. Future studies should assess the acute effects of low-dose PFOS on insulin secretion *in vivo*, and whether repeated or continuous exposure to PFOS eventually leads to impaired insulin secretion.

Given the structural similarities between PFOS and long-chain fatty acids, previous studies proposed that the effects of PFOS on insulin secretion are mediated by GPR40 [20,21]. Specifically, co-treating beta-TC-6 cells with PFOS and GW1100 (a GPR40 antagonist) attenuated the PFOS-induced hypersecretion of insulin under LG conditions [21]. Additionally, exposure to 10 mg/kg PFOS *in vivo* hyperstimulated insulin secretion in wild-type mice but not in GPR40 knockout male mice 1-hr post-injection [21]. Surprisingly, in our current study, we did not see evidence that GPR40 was mediating the acute stimulatory effects of PFOS on insulin secretion in female mouse islets *in vitro*. In fact, the effect of PFOS on insulin secretion was amplified in the presence of GW1100. The discrepancy between our study and previous *in vitro* studies may be because our study was conducted in primary mouse islets rather than immortalized beta cells. Additionally, given that we observed PFOS-induced hypersecretion of insulin under HG conditions in our study (both *in vitro* and *in vivo*), we only administered PFOS ± GW1100 under HG rather than LG conditions. In doing so, we simultaneously triggered both a GSIS response and paracrine signaling. Lastly, our exposure concentrations for both PFOS and GW1100 differed from previous studies. Although we do not report a role of GPR40 in our study, we cannot rule out the possibility that the effects of PFOS are partially mediated by other G-protein coupled receptors (GPCRs), such as GPR120, which also binds fatty acids.

Since PFOS increased GSIS in mouse islets but decreased GSIS in INS-1 832/3 cells, we hypothesized that other endocrine cell types may be mediating the observed phenotype in mouse islets. Therefore, we assessed the involvement of GLP-1R, a receptor on beta cells that amplifies GSIS when bound by glucagon or GLP-1 secreted from the alpha cells [29]. Interestingly, we report that co-exposure of PFOS with the GLP-1R antagonist Ex(9-39) partially mitigated the effects of PFOS on insulin secretion in islets; however, PFOS exposure had no effect on glucagon secretion under HG conditions. These data suggest that PFOS partially amplifies GSIS by stimulating GLP-1 secretion from the alpha cell, but this remains to be determined. Additionally, although our Ex(9-39) dose aligned with previous studies [45], we cannot rule out the possibilities that this dose was not sufficient to block all GLP-1 receptors on the beta cell, since we did not fully ablate the PFOS-induced amplification of GSIS. Alternatively, these data may suggest that other signalling pathways may be involved in this phenotype. For example, there is evidence to suggest that PFAS interacts with peroxisome proliferator-activated receptors (PPARs) in bone and adipose tissue [46,47]. PPARs are receptors that are highly expressed in several cell types [48], including beta cells, and are known to enhance GSIS and regulate fatty acid metabolism [49,50]. Future studies should continue to investigate the impact of PFOS on intra-islet signalling pathways, including GLP-1, GPCRs, and PPARs.

This is also the first study to evaluate the effects of PFOS exposure on primary human donor islets. These data are more difficult to interpret since we did not observe reproducible effects of PFOS on insulin secretion. This could be due to inherent biological and environmental variability and/or differences in islet isolation and culture [51,52]. Congruent with the data in primary mouse islets, we observed a stimulation of GSIS in donors R430 and R471. However, we also observed a suppression of GSIS in R496 and no effect on GSIS in R447 following acute PFOS exposure. Prolonged PFOS exposure had no effect on GSIS in donors R430, R447, and R471 but stimulated GSIS In R496. Although these findings do not align with our data in INS-1 832/3 cell or mouse islets, the prolonged PFOS data for donor R496 aligns with previous findings showing a stimulation in GSIS in EndoC-betaH1 cells after a 72-hr exposure to 100 nM – 1 µM PFOS [25]. Collectively, our findings suggest that PFOS exposure can dysregulate insulin secretion in primary human donor islets, although future studies in a larger sample size are warranted to better understand the heterogeneity across donors.

Our study includes the largest report of PFAS measurements in human plasma and pancreas to date. To our knowledge, this is also the first study to investigate the effects of PFOS exposure on insulin secretion across several model systems, including immortalized beta cells, mouse islets and human donor islets. Collectively, we show that PFOS preferentially accumulated in human and mouse pancreas, and acutely stimulates GSIS, pointing to PFOS as an islet toxicant. We also report that the acute effects of PFOS on insulin secretion varies between models, with mouse islets showing the most robust and reproducible findings. This emphasizes the importance of carefully selecting model systems when conducting toxicological assessments.

## 5. ACKNOWLEDGEMENTS

This research was supported by a Canadian Institutes of Health Research (CIHR) Project Grant (#PJT-186282). J.E.B. is supported by a Dorothy Killam Fellowship and an Early Researcher Award from the Ontario Government. J.P. was supported by the Guiding interdisciplinary Research on Women’s and girls’ health and Wellbeing (GROWW) scholarship and Ontario Graduate Scholarship. M.P.H. was supported by a CIHR CGS-D award. E.v.Z. was supported by an NSERC CIRTN-R2FIC-CREATE award. L.B. was supported by an NSERC CIRTN-R2FIC-CREATE and CIHR CGS-D award. M.E.A.C. was supported by an NSERC CGS-M and NSERC CGS-D award.

This work includes data from HumanIslets.com funded by the Canadian Institutes of Health Research, JDRF Canada, and Diabetes Canada (5-SRA-2021-1149-S-B/TG 179092) with data from islets isolated by the Alberta Diabetes Institute IsletCore with the support of the Human Organ Procurement and Exchange program, Trillium Gift of Life Network, BC Transplant, Quebec Transplant, and other Canadian organ procurement organizations.

## 6. AUTHOR CONTRIBUTIONS

J.P., M.P.H. and J.E.B. conceived the experimental design, conducted the data analysis, and wrote the manuscript. J.P., M.P.H., L.B., M.E.A.C., A.A., A.B., A.T., A.A.R. and J.E.B. were involved with data acquisition and interpretation of data. All authors contributed to manuscript revisions and approved the final version of the article.

## ABBREVIATIONS

GLP-1: Glucagon-like peptide-1
GPR40: G-protein coupled receptor-40
GSIS: Glucose-stimulated insulin secretion
HOMA-beta: Homeostatic model assessment of beta cell function
HOMA-IR: Homeostatic model assessment of insulin resistance
PFAS: Poly- and perfluoroalkyl substances
PFHxS: Perfluorohexanesulfonic acid
PFOA: Perfluorooctanoic acid
PFOS: Perflurooctane sulfonic acid
POPs: Persistent organic pollutants

