## Supplemental figures for "Acute PFOS exposure consistently dysregulates glucose-stimulated insulin secretion across model systems"

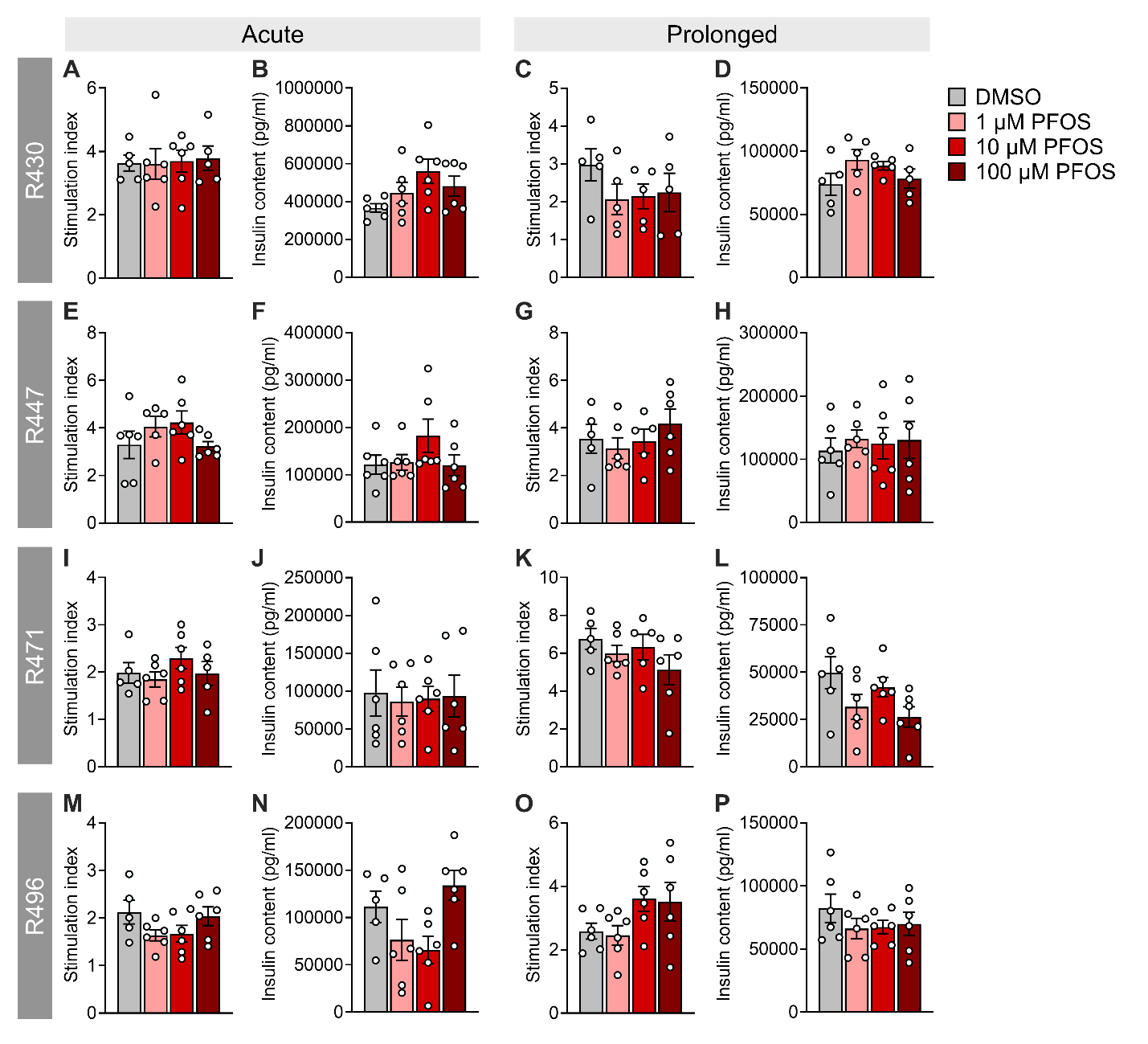

**Supplemental Figure 1: Insulin content and stimulation index following acute and prolonged GSIS assays in human donor islets.** Islets from 4 human organ donors were exposed to DMSO (vehicle) or PFOS (1, 10, or 100 µM) either acutely during a GSIS assay or for 48-hrs and islet function was subsequently assessed with a static GSIS (see **Figure 4A,B** for experimental design schematic). Stimulation index following **(A,E,I,M)** acute or **(C,G,K,O)** prolonged PFOS exposure; stimulation index was calculated as a ratio of insulin concentration under HG relative to LG (raw data is presented in **Figure 4**). Insulin content following **(B,F,J,N)** acute or **(D,H,L,P)** prolonged PFOS exposure. Data is presented as mean ± SEM. Individual data points represent technical replicates from a single donor (n = 4–6/technical replicates/donor/condition). The following statistical tests were used: one-way ANOVA with Dunnett’s post-hoc.
